# Knocking out *AtCALS7* disturbs photoassimilate distribution and weakens defence against phytoplasma infection

**DOI:** 10.1101/2021.06.25.449948

**Authors:** Chiara Bernardini, Amit Levy, Sara Buoso, Alberto Loschi, Simonetta Santi, Marta Martini, Federico Bosetto, Stacy Welker, Joon Hyuk Suh, Yu Wang, Christopher Vincent, Myrtho O’Pierre, Aart J. E. van Bel, Rita Musetti

## Abstract

Callose accumulation around sieve pores, under control of Callose synthase 7 (AtCALS7), has been interpreted as a mechanical response to limit pathogen spread in phytoplasma-infected plants. AtCALS7 is also involved in sieve-pore development and, hence, in mass-flow regulation, carbohydrate metabolism and distribution, and plant growth. Multiple roles of AtCALS7 during phytoplasma infection were investigated in healthy and phytoplasma-infected [Chrysanthemum Yellows (CY)-phytoplasma] wild-type and At*cals7ko* Arabidopsis lines. In keeping with an increased phytoplasma titre in *Atcals7ko* plants, floral stalk height of infected wild-type and mutant plants was reduced by, respectively, 88 and 100% in comparison to their healthy controls, suggesting a higher investment of host resources in phytoplasma growth in the absence of AtCALS7. The apparently increased susceptibility of mutants was investigated by microscopic, metabolic and molecular analyses. Infection influenced the sieve-pore functionality in wild-type plants, which hardly affected plant growth, and plasmodesmata in the cortex, a phenomenon less prominent in mutants. Infection also increased the level of some sugars (glucose, sucrose, myoinositol), but to the highest extent in mutants. Finally, infection induced a similar upregulation of gene expression of enzymes involved in sucrose cleavage (*AtSUS5*, *AtSUS6*) in mutants and wild-type plants and an upregulation of carbohydrate transmembrane transporters (*AtSWEET11*, *AtSTP13*, *AtSUC3*) in mutants only. A more effective plasmodesmal closure seemingly suppressed spread of phytoplasma effectors, which rendered wild-type plants less susceptible to infection, because gene expression of enzymes channeling carbohydrates towards phytoplasmas is less promoted.

**One sentence summary:** Knocking out AtCALS7 disturbs photoassimilate distribution between axial and terminal sinks and weakens the defence against phytoplasma infection.

## Introduction

Phytoplasmas are wall-less, pleomorphic plant pathogens belonging to the class *Mollicutes*. They are confined to the sieve elements (SEs) of host plants (van Bel and Musetti, 2019) or to the body of phloem-feeding insect vectors (Alma et al., 2019). Phytoplasmas cause serious yield losses and affect the quality of crops of economic interest (Albertazzi et al., 2009). Profound alterations in transcriptome and proteome (Albertazzi et al., 2009; Cao et al., 2017; 2019), and in the phytohormone balance of infected plants (Dermastia, 2019; Bernardini et al., 2020) are reflected by a vast range of symptoms, such as witches’-brooms, leaf chlorosis, virescence, phyllody, and floral abortion (Ermacora and Osler, 2019), often leading to sterility and unproductiveness of the host plant (Namba et al., 2019).

Plants react to phytoplasma invasion by mechanical occlusion of sieve pores, which is due to a fast plugging by specialized proteins (Will and van Bel, 2006; Furch et al., 2007; Pagliari et al., 2017), followed by a slower constriction due to additional deposition of callose around the pores (Musetti et al., 2010; Santi et al., 2013a; Musetti et al., 2013). In *Arabidopsis thaliana*, 12 genes (*CALS1*-*12*) encoding for callose synthase enzymes have been found (Xie and Hong, 2011). The diverse *CALS*-gene expression patterns throughout the plant are indicative of different local roles (Ellinger and Voigt, 2014). As for the phloem, Barratt et al., (2011) and Xie et al., (2011) demonstrated that the *AtCALS7* gene encodes for Callose Synthase 7 (AtCALS7), the enzyme responsible for callose deposition around sieve pores. AtCALS7 also regulates sieve-plate development during phloem differentiation, which is an important determinant of mass flow in mature sieve tubes, and, hence, influences sink carbohydrate availability and plant growth (Xie et al., 2011).

Synthesis of the cell-wall polymer callose (1,3-β-D-glucan) requires sucrose units that are cleaved by sucrose synthases into fructose and UDP-glucose, the substrate for callose synthase. Arabidopsis callose synthase proteins contain multiple transmembrane domains that are clustered into two regions (N-terminal and C-terminal), leaving a large hydrophilic central loop that faces the cytoplasm. This loop contains the putative catalytic domain, which has been further subdivided into two domains: the UDP-glucose binding domain and the glycosyltransferase domain (Chen and Kim, 2009). In the sieve tubes of Arabidopsis, two membrane-bound sucrose synthase isozymes, AtSUS5 and AtSUS6 (Barratt et al., 2009), associated with AtCALS7 form a unique enzyme complex (Bieniawska et al., 2007; Ruan, 2014; Stein and Granot, 2019).

Sucrose is of paramount importance in plant-pathogen interactions (Bolton et al., 2009; Rojas et al., 2014; Fatima and Senthil-Kumar, 2015; Dodds and Lagudah, 2016; Lee et al., 2016). Sugars are involved in several signaling processes (Lecourieux et al., 2014) and contribute to the plant immune response, as priming molecules for rapid activation of defense against biotic and abiotic stresses (Bolouri-Moghaddam and van Den Ende, 2012). Moreover, sucrose is the ideal carbon source owing to its high concentration in SEs (van Bel and Musetti, 2019). As result, sucrose host metabolism is often affected by phytoplasmas (Santi et al., 2013b; De Marco et al., 2016; Prezelj et al., 2016) given their full dependence on the host (Oshima et al., 2013; Musetti et al., 2016).

Assumed the intricate and interdependent roles of callose and sugars in phytoplasma infection and the functional connection between callose synthase and sucrose synthases (Bieniawska et al., 2007; Ruan, 2014; Santi et al., 2013a; Stein and Granot, 2019) we investigated the consequences of phytoplasma infection and the role of CALS7 in sucrose metabolism and cell-cell transport of carbohydrates. Wild-type and *Atcals7ko* [a mutant line in which the gene encoding for AtCALS7 was silenced (Barratt et al., 2011)] Arabidopsis plants, healthy or infected with Chrysanthemum Yellows (CY)-phytoplasma, were compared regarding plant morphology, sieve-tube substructure, gene expression and carbohydrate amount. We investigated whether the loss of AtCALS7 in CY-infected Arabidopsis would affect plant growth, symptom expression and phytoplasma titre. Furthermore, we evaluated whether the loss of AtCALS7 had an impact on: i) the SE ultrastructure; ii) the distribution of callose deposits at the plasmodesmata in midrib cortical parenchyma and in leaf epidermis; iii) the amount of sucrose, glucose, fructose, myoinositol, sorbitol, arabinose and raffinose in midrib tissues; iv) the expression patterns in midrib tissues of several genes involved in sugar transport and metabolism.

The results indicate that, apart from the negative role in the sieve-plate occlusion, loss of AtCAL7 appears to confer increased susceptibility to CY-phytoplasma infection, due to alterations in symplasmic connectivity and expression of genes involved in sugar metabolism and membrane transport in *Atcals7ko* plants.

## Results

### Phloem translocation speed

The speed of phloem translocation was measured in healthy wild-type and *Atcals7ko* Arabidopsis plants (Fig. 1) through the label with C^14^. The linear translocation velocity (expressed as cm h^-1^) was approximatively 50% lower in *Atcals7ko* mutants than in wild-type plants. The average speed in wild-type plants was 10.2 ± 1.6 cm h^-1^, while it was 5.0± 2.0 cm h^-1^ in mutants. As the infected plants developed very short stems (wild-type) or failed to do so (*Atcals7ko*, Fig. 2C), it was not possible to determine the translocation speed in CY-infected plants.

**Fig. 1.**
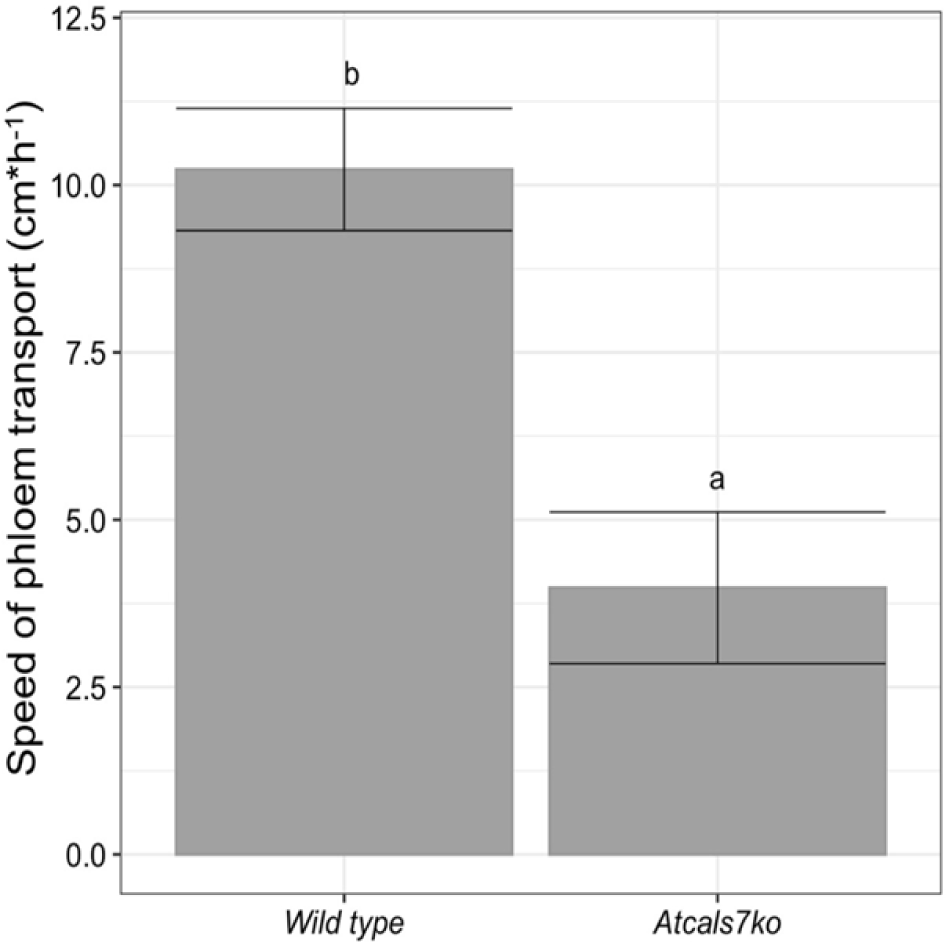
Phloem transport speed in wild-type and *Atcals7ko* Arabidopsis lines. Carbohydrate translocation speed along the phloem, measured with ^14^C isotope. The speed is expressed as cm/h and it is calculated by average time of arrival of the ^14^C label in the stem phloem tissue near the x-ray detector. Statistical analysis was performed using the Tukey HSD test as the *post hoc* test in an one-way ANOVA. Different letters (a, b) above the bars indicate significant differences, with P < 0.05. Error bars indicate the Standard Error of the Mean of 4 biological replicates for each condition.

**Fig. 2.**
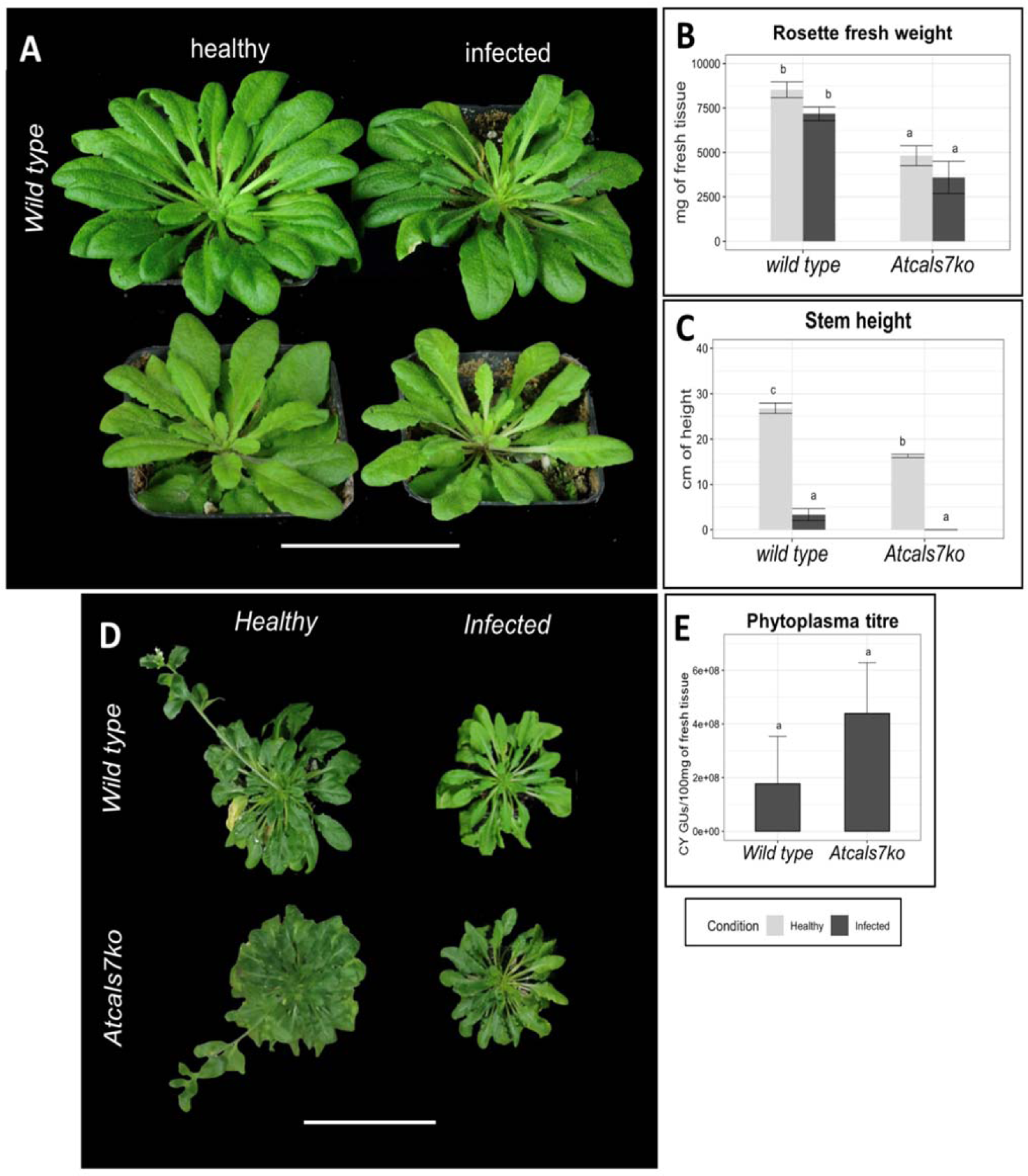
Plant phenotype and phytoplasma titre in wild-type and *Atcals7ko* line. Representative images of healthy and CY-infected wild-type and *Atcals7ko* plants. Following CY-infection, at 20 days after the inoculation access period (IAP), both plant groups showed yellowish small leaves. Leaves emerged after phytoplasma inoculation were shorter, with thicker main vein and smaller petiole (**A**). Regardless phytoplasma infection, rosette fresh weight is significantly reduced in *Atcals7ko* plants in comparison with wild-type (**B**). In 60-day-old Arabidopsis plants, grown under long day conditions, stem was well developed in healthy wild-type but reduced in length in *Atcals7ko* plants (**C, D**). Following CY infection, stem length resulted strongly reduced in wild-type individuals and absent in the *Atcals7ko* mutants (**C**, **D**). Although not significantly different, phytoplasma titre increases in the mutant line in comparison to the wild type (**E**). Different letters indicate different means according to the non-parametric Kruskal-Wallis *post hoc* test, P<0.05. Error bars indicate Standard Error of the Mean of 8 biological replicates for each condition. Plant weight (**B**) 20 days IAP expressed as mg of fresh tissue. Statistical analysis was performed using the Tukey HSD test as the *post hoc* test in a two-way ANOVA. Different letters (a, b) above the bars indicate significant differences, with P < 0.05. Error bars indicate the Standard Error of the Mean of 8 biological replicates for each condition.

### Phenotypes of healthy and CY-infected Arabidopsis lines and phytoplasma titre

Symptom appearance and rosette growth and weight (Fig. 2A and B) of wild-type and *Atcals7ko* plants, healthy or infected by CY-phytoplasmas were compared. Floral stem length was also evaluated in plants grown under long-day light conditions (Fig. 2C). Moreover, phytoplasma titre was quantified in infected Arabidopsis lines by qPCR (Fig. 2E).

*Atcals7ko* plants were smaller than wild-type plants, having stunted growth (Fig. 2A) and leaves with a thick main vein. Fresh weight of rosettes (Fig. 2B) was reduced in *Atcals7ko* plants (average weight: 4.8 ± 1.5 g) to the 56% as compared to the wild-type plants (average weight: 8.5 ± 1.2 g). Following CY-infection, rosette fresh weight, on average, was affected by a reduction of 16% in wild-type plants (7.1 ± 1.0 g) and 27% in the *Atcals7ko* line (3.5 ± 2.2 g) in comparison with their respective healthy controls. In both lines, leaves that emerged after phytoplasma inoculation were yellowish, small, narrowed, with a thicker main vein and a shorter petiole. When grown under long-day light conditions, the mutant line developed a floral stalk reduced to the 41% in length (average length: 16.3 ± 0.7 cm) in comparison to the wild-type (average length 26.8 ± 2.5 cm) (Fig. 2C and D). CY-infected wild-type plants produced a stalk reduced by 88% (average length 3.3 ± 3.2 cm) compared to the healthy controls, while CY-infected *Atcals7ko* mutants failed to produce a stalk at all (Fig. 2C and D). Although not significantly different, phytoplasma titre resulted, on average, higher in mutant plants (i.e. 4.4E+08 phytoplasma GUs in 100 mg of leaf sample) than in the wild-type (i. e. 1.8E+08 phytoplasma GUs in 100 mg of leaf sample) (Fig. 2E).

### Electron-microscopic observations on midrib vascular bundles

To visualize changes in SE ultrastructure due to mutation effects and/or pathogen presence, ultrathin sections of midrib vascular bundles were examined under a transmission electron microscope (TEM). For every condition, five non-serial sections from 5 different plants of both lines were analysed. Healthy wild-type samples showed a regular SE and CC ultrastructure (Fig. 3A-D). In lateral (Fig. 3B) and transverse (Fig. 3C) sieve plates, sieve pores were surrounded by a callose lining (Fig. 3B and C) that did not constrict their lumen. At the SE/CC wall interface, sieve endoplasmic reticulum (SER) was visible in front of the open, typically branched pore-plasmodesma units (PPUs, Fig. 3D). In infected wild-type midribs (Fig. 3E-H), numerous phytoplasmas were visible in the SE lumen (Fig. 3E) or in proximity of the sieve plates (Fig. 3F). In transverse sieve plates (Fig. 3G), the pores were constricted by callose deposition (Fig. 3G), whereas they appeared mostly open in lateral sieve plates (Fig. 3F). PPUs were open and displayed a slightly thickened callose lining at the SE side (Fig. 3H). As in healthy plants, SER was observed near the PPU opening in infected plants (Fig. 3H).

**Fig. 3.**
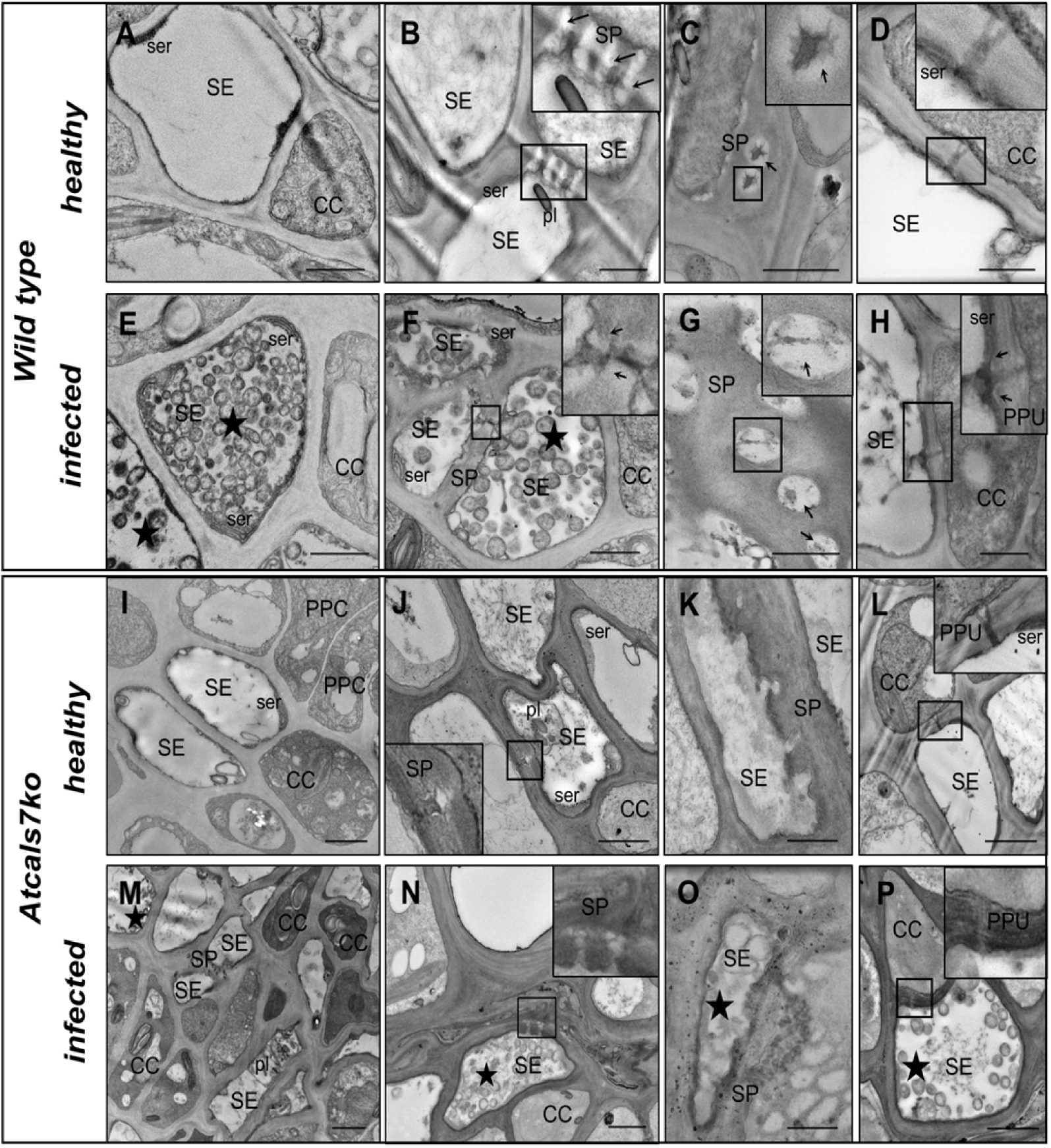
Representative TEM micrographs of the sieve elements of healthy and CY-infected Arabidopsis lines. **A-H.** Cross-sections of midribs from healthy (**A-D**) and infected (**E-H**) wild-type Arabidopsis leaves. Healthy samples present unaltered sieve elements and companion cells (**A**), with a regular shape and no signs of necrosis or subcellular aberrations. The sieve pores, at both lateral (**B**) and ordinary (**C**) sieve plates, are not constricted by callose collars, pore-plasmodesma units are open and show their typical branched shape with sieve endoplasmic reticulum located in the proximity of the orifice (**D**). In infected midribs, numerous phytoplasmas are visible inside the sieve elements (**E)** and at the sieve plate (**F**). Lateral sieve plates have sieve pores mostly open (**F**), whereas in ordinary sieve plates (**G**) pores are constricted by callose depositions. Pore-plasmodesma units display a similar morphology as in healthy samples, with a thin callose line at the sieve-element side (**H**). **I-P**. Cross-sections of midribs from healthy (**I-L**) and CY-infected (**M-P**) *Atcals7ko* Arabidopsis leaves. In healthy *Atcals7ko* samples, phloem cells are apparently well structured (**I**), but the sieve plates show aberrant morphology (**J**, **K**). Pores lack callose and appear not fully developed (**J**, **K**). Pore-plasmodesma units are branched, similar as in wild-type samples (**L**). In CY-infected *Atcals7ko* samples, many phloem cells evidence thick cell walls (**M**), others are collapsed (**N**). Sieve plates are deformed, thickened (**N**, **O**) and pores are filled by electron-opaque material (**N**). Pore- plasmodesma units appeared large, without well-defined branches (**P**). Arrows: callose; CC: companion cell; PPC: phloem parenchyma cell; pl: plastid; PPU: pore-plasmodesma unit; SE: sieve element; ser: sieve element reticulum; SP: sieve plate; star: phytoplasmas. Bars correspond to 1μm.

In the healthy *Atcals7ko* line (Fig. 3I-L), SEs and CCs appeared identical to those in the wild-type (Fig. 3I). Sieve-element protein filaments, plastids and SER were readily recognizable in the SE lumen or close to the plasma membrane (Fig. 3J). In accordance with previous studies (Barratt et al., 2011; Xie et al., 2011), however, sieve plates lacked callose and showed an aberrant morphology (Fig. 3J and K). Some sieve-pore channels seemed to be incompletely open or not fully developed (Fig. 3J and K), whereas PPUs displayed a normal, branched appearance (Fig. 3L). Inside SEs of the infected *Atcals7ko* line phytoplasmas were visible (Fig. 3M-P). Many SEs possessed thick walls (Fig. 3M), while others had collapsed (Fig. 3N). Like in healthy samples, sieve plates appeared to be damaged, with sieve pores filled by electron-opaque material (Fig. 3O). PPUs were enlarged without a well-defined branching (Fig. 3P). In the proximity of PPUs, SER was recognizable in the SEs of both healthy and CY-infected plants (Fig. 3L and P).

### Confocal laser scanning microscopy analyses and imaging

Confocal laser scanning microscopy (CLSM) in combination with aniline blue staining were used to evaluate presence, number and distribution of blue-fluorescent callose deposits, the fluorescence intensity and the integrated density (i.e. the sum of intensity within the area of the region of interest) in various leaf tissues of wild-type and *Atcals7ko* plants, healthy or CY-infected. Fluorescent regions represented the callose lining along the symplasmic corridors of SE/CC complexes (Fig. 4); the discrete fluorescent dots in midrib cortical parenchyma (Fig. 4) and epidermis (Fig. 5) localized callose collars around plasmodesmata. In infected wild-type samples, intense aniline blue signals revealed a significant increase in callose deposits at the SE level (Fig. 4B), in comparison to their respective healthy controls (Fig. 4A). The level of fluorescence (Fig. 4E), the number (Fig. 4F) and the integrated density (i.e. the sum of intensity within the area of the region of interest) of deposits (Fig. 4G) were measured and compared in healthy and infected wild- type samples. Infection induced a significant increase in callose deposition in SEs of wild- type plants (Fig. 4E-G). As expected, no fluorescence was detected in the corresponding SE area in *Atcals7ko* samples, even in case of phytoplasma infection (Fig. 4C and D).

**Fig. 4.**
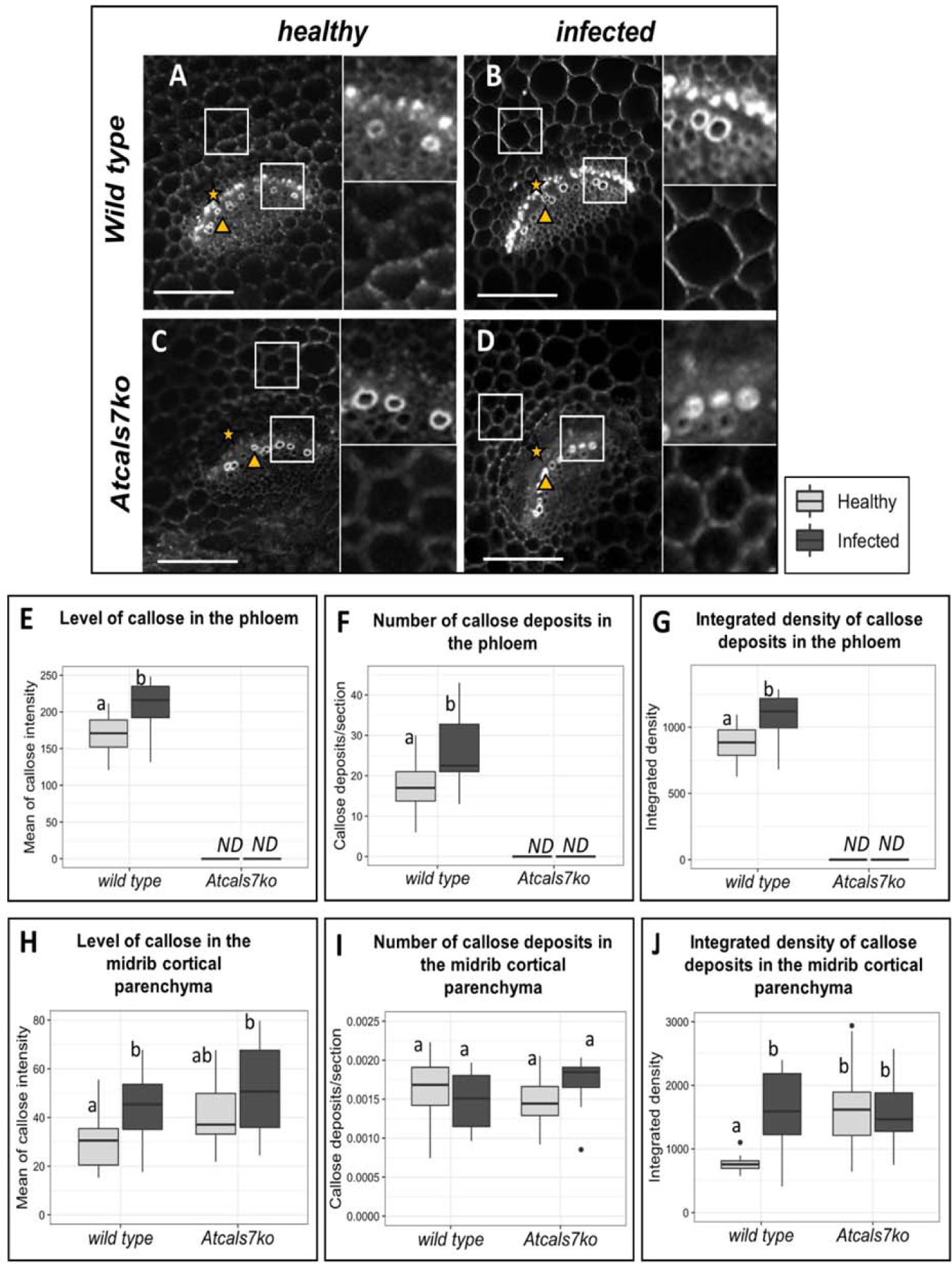
Single-plane confocal micrographs and imaging of callose deposits at plasmodesmata in phloem area and in parenchyma of healthy and CY-infected Arabidopsis midribs. Fluorescent spots indicate callose depositions in the phloem area of healthy (**A**) and CY-infected (**B**) wild-type samples. The fluorescence intensity (**E**), the number (**F**) and the integrated density [i.e. the sum of the intensity in the region of interest (ROI)] of the deposits (**G**) is significantly higher in the infected midribs compared to the healthy ones. No signal is visible in *Atcals7ko* midribs (**C**, **D**). Punctate dots, indicating callose deposits at plasmodesmata are visible in the midrib parenchyma of all samples (**A**-**D**). In the ROI, fluorescence intensity (**H**) and number of dots (**I**) are not different in healthy samples, but the integrated density (i.e. the sum of the intensity within the area of the region of interest, ROI) is significantly higher in the mutant line than in the wild-type (**J**). Following infection, the fluorescence intensity and the integrated density of the signal in the ROI significantly increased only in wild-type samples. In the pictures, stars indicate phloem and triangle indicate xylem. Statistical analysis was performed using the Tukey HSD test as the *post hoc* test in a two- way ANOVA. Different letters (a, b, c, d) above the bars indicate significant differences, with P < 0.05. Boxplots were obtained from three biological replicates for each condition and almost 10 technical replicates for each sample. Bars correspond to 100μm.

**Fig. 5.**
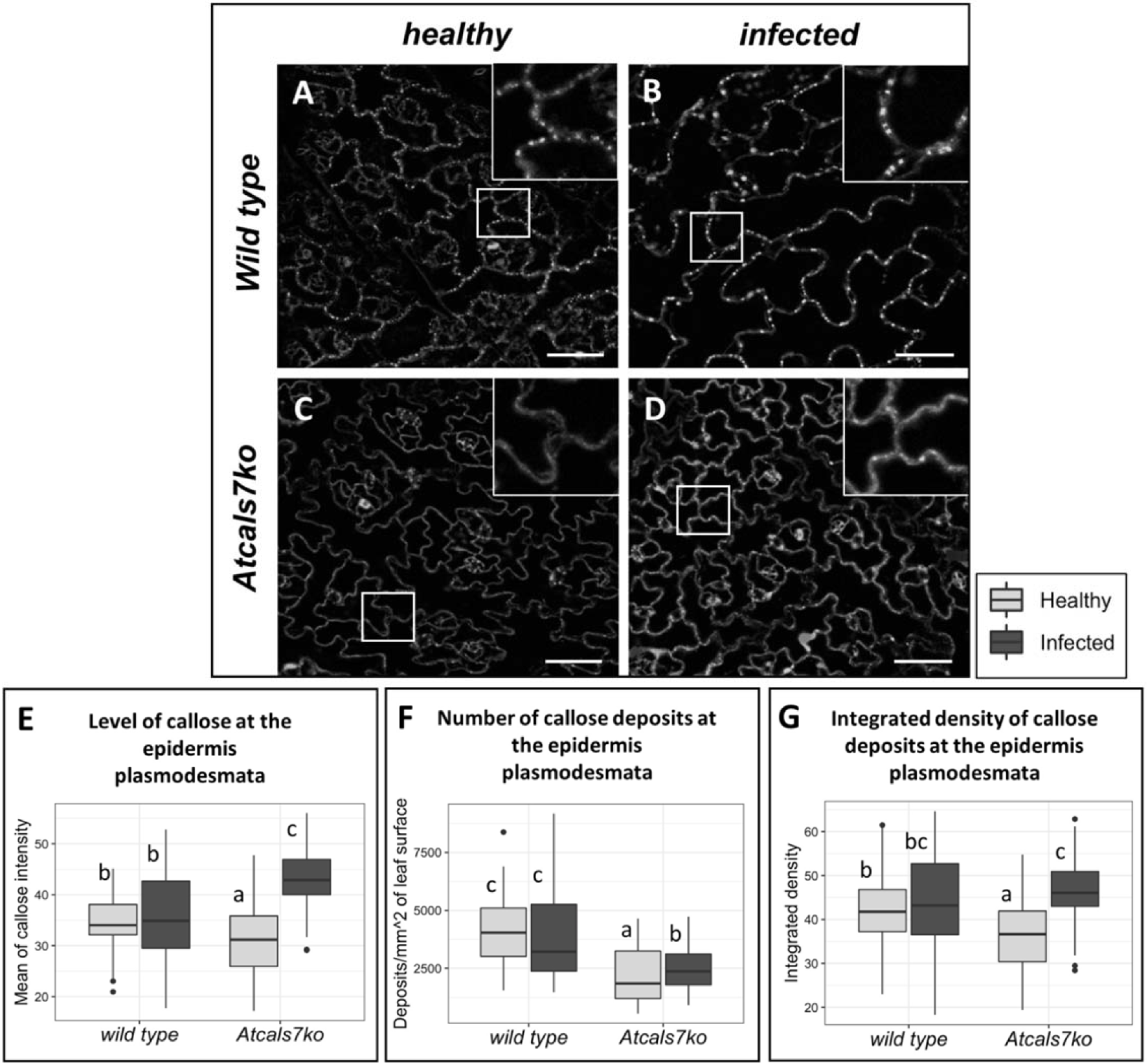
Single-plane confocal micrographs and imaging of callose deposits at plasmodesmata in epidermal cells. Aniline blue fluorescent dots (**A**-**D**), their intensity (**E**), number (**F**) and integrated density (i.e. the sum of the intensity within the area of the region of interest, ROI, **G**) were assayed at leaf epidermis level in healthy and infected samples of both lines. In healthy samples, the intensity (**E**) and the number of fluorescent dots (**F**), and the integrated density (**G**) are lower in *Atcals7ko* line compared to the wild-type. Following CY-infection the three parameters significantly increased only in *Atcals7ko* line. Statistical analysis was performed using the Tukey HSD test as the *post hoc* test in a two-way ANOVA. Different letters (a, b, c) above the bars indicate significant differences, with P < 0.05. Boxplots were obtained from three biological replicates for each condition and almost 10 technical replicates for each sample. Bar corresponds to 50μm.

Image analysis allowed us to quantify the fluorescent signals (plasmodesma-located dots) in cell walls of midrib cortical parenchyma (Fig. 4). In healthy samples of both Arabidopsis lines, the signal intensity (Fig. 4H) and number of fluorescent dots (Fig. 4I) did not differ significantly in the selected areas. However, the integrated density was significantly higher in mutant line than in wild-type plants. Following infection, the level of fluorescence and the integrated density significantly increased only in wild-type samples and not in the *Atcals7ko* line (Fig. 4H-J). In contrast to the situation in the cortical parenchyma, the intensity (Fig. 5E), the number of fluorescent dots (Fig. 5F), and the integrated density (Fig. 5G) in the epidermis of healthy plants were lower in *Atcals7ko* than in wild-type samples. Following CY-infection, the three parameters investigated increased significantly only in *Atcals7ko* line. In particular, the signal intensity in the infected mutant line was the highest as compared to the other samples (Fig. 5E), whereas the integrated density was similar to that found in infected wild-type samples (Fig. 5G). With exception of autofluorescence of the xylem and chloroplasts no fluorescence signal was detected in unstained samples (Supplementary file 1).

### Sugar quantification in midrib tissues

Sucrose, glucose, fructose, myoinositol, sorbitol, arabinose and raffinose were quantified in the midribs of wild-type and *Atcals7ko* plants (Fig. 6). No significant differences between both lines were found in absence of infection (Fig. 6A-G). Following CY-infection, the amounts of the above-mentioned sugars did not differ significantly from the controls in the wild-type line (Fig. 6A-G). In the *Atcals7ko* line, sucrose, glucose and myoinositol significantly increased in response to CY-infection (Fig. 6A, B and D). In particular, comparing with the amounts found in the midribs of healthy *Atcals7ko* plants, sucrose increased around 5-fold in infected *Atcals7ko* plants (Fig. 6A), glucose around 4- fold (Fig. 6B), myoinositol around 2-fold (Fig. 6D).

**Fig. 6.**
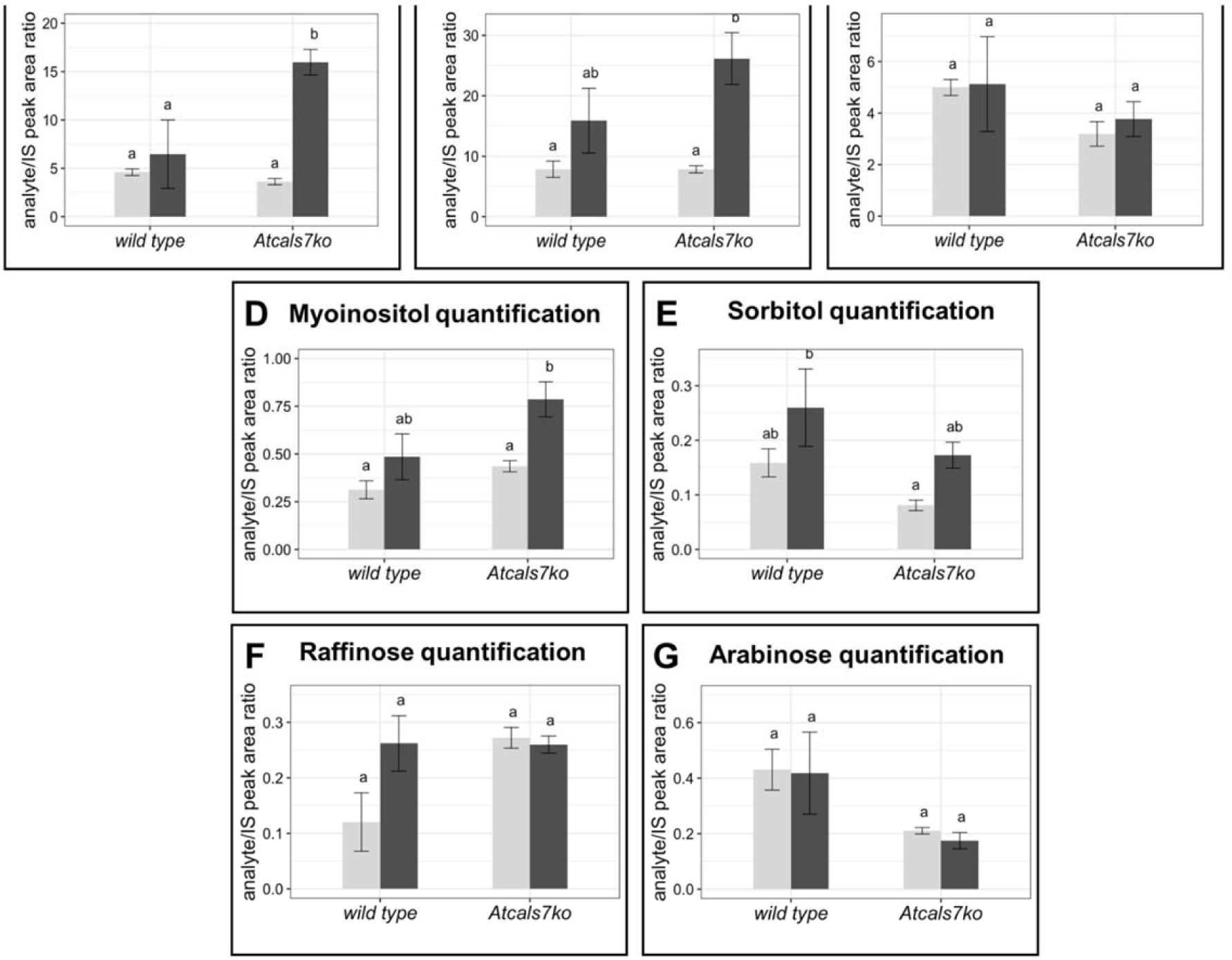
Sugar quantification in the midribs of healthy and infected Arabidopsis lines. Sucrose (**A**), glucose (**B**), fructose (**C**), myoinositol (**D**), sorbitol (**E**), raffinose (**F**) and arabinose (**G**) were quantified in midribs of the two different lines, healthy or CY-infected. Data obtained were expressed as analyte/IS peak area ratio. Statistical analysis was performed using the Tukey HSD test as the *post hoc* test in a two-way ANOVA. Different letters (a, b) above the bars indicate significant differences, with P < 0.05. Error bars indicate the Standard Error of the Mean of 4 biological replicates for each condition run in triplicate.

### Gene expression analyses

The expression level of the sieve-element specific callose synthase 7 gene (*AtCALS7*) was analyzed in midribs of healthy and CY-infected wild-type plants and was significantly up-regulated (around 2.5-fold) in infected plants (Fig. 7A). Moreover, the expression of genes involved in sugar metabolism and transport, in the midribs of source leaves of the Arabidopsis lines was investigated. Expression levels of sucrose synthases (*AtSUS5* and *AtSUS6*), sucrose transporters (*AtSUC2* and *AtSUC3*), sugar transport facilitators (*AtSWEET11, AtSWEET12*), cell wall invertases (*AtCWINV1*, *AtCWINV6*) were determined in the four plant groups under investigation (Fig. 8B-F).

**Fig. 7.**
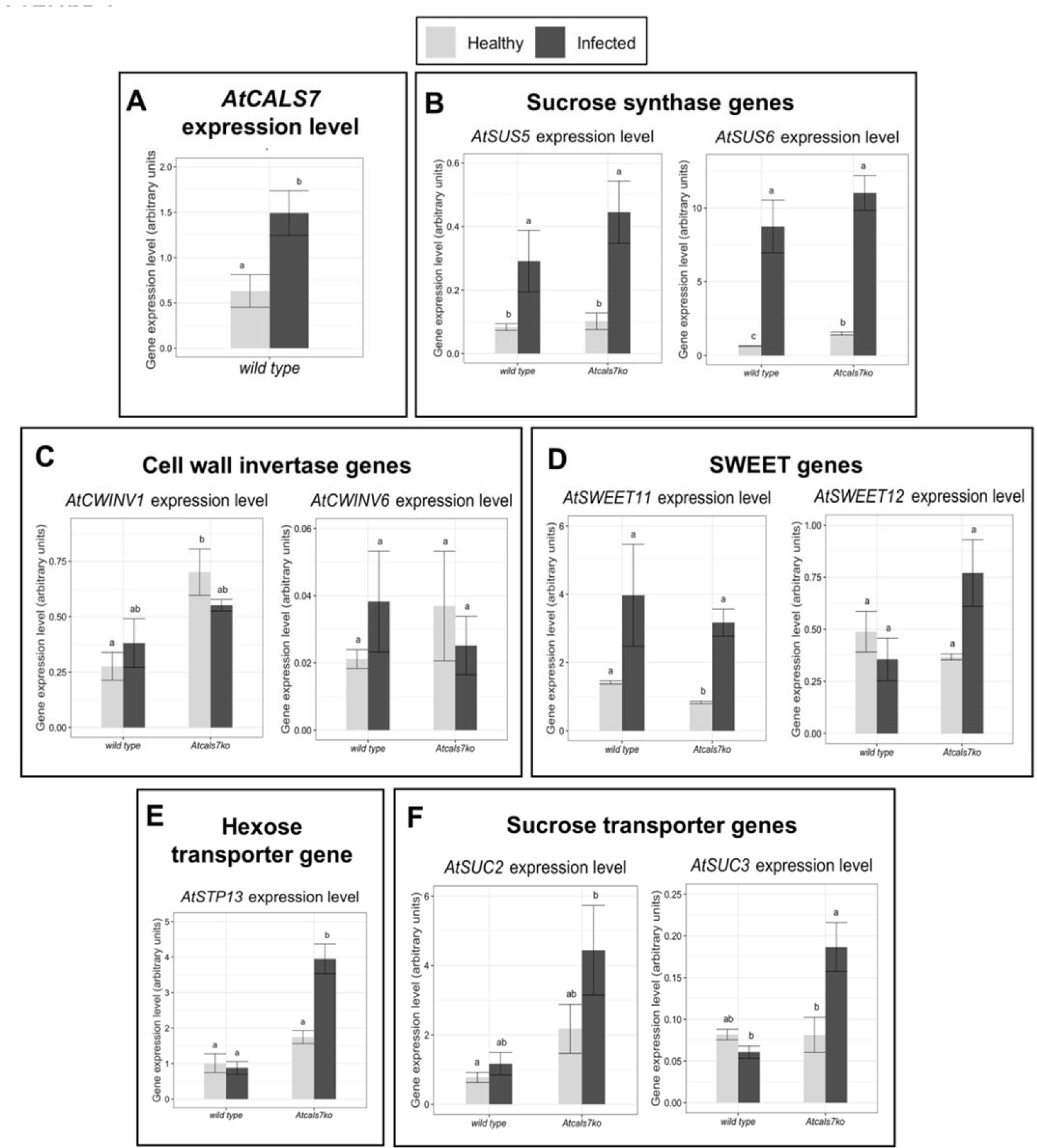
Transcript profiling of genes involved in phloem callose synthesis and sugar transport and metabolism. **A**. Expression level of phloem callose synthase gene (*AtCALS7*) in healthy and CY-infected wild-type plants. **B**-**F**. Transcript profiling of sucrose synthases AtSUS5 and AtSUS6 (**B**), sucrose transporters AtSUC2 and AtSUC3 (**C**), SWEET sugar facilitators AtSWEET11 and AtSWEET12 (**D**), cell wall invertases AtCWINV1 and AtCWINV6 (**E**), the hexose transporter AtSTP13 (**F**). Healthy and infected plants belonging to the two lines were compared. Expression values were normalized to the *UBC9* transcript level, arbitrarily fixed at 100, then expressed as mean normalized expression ±SD (transcript abundance). Statistical analysis was performed using the Tukey HSD test as the *post hoc* test in a two-way ANOVA. Different letters (a, b, c, d) above the bars indicate significant differences, with P < 0.05. Error bars indicate the Standard Error of the Mean of 5 biological replicates for each condition run in triplicate.

**Fig. 8.**
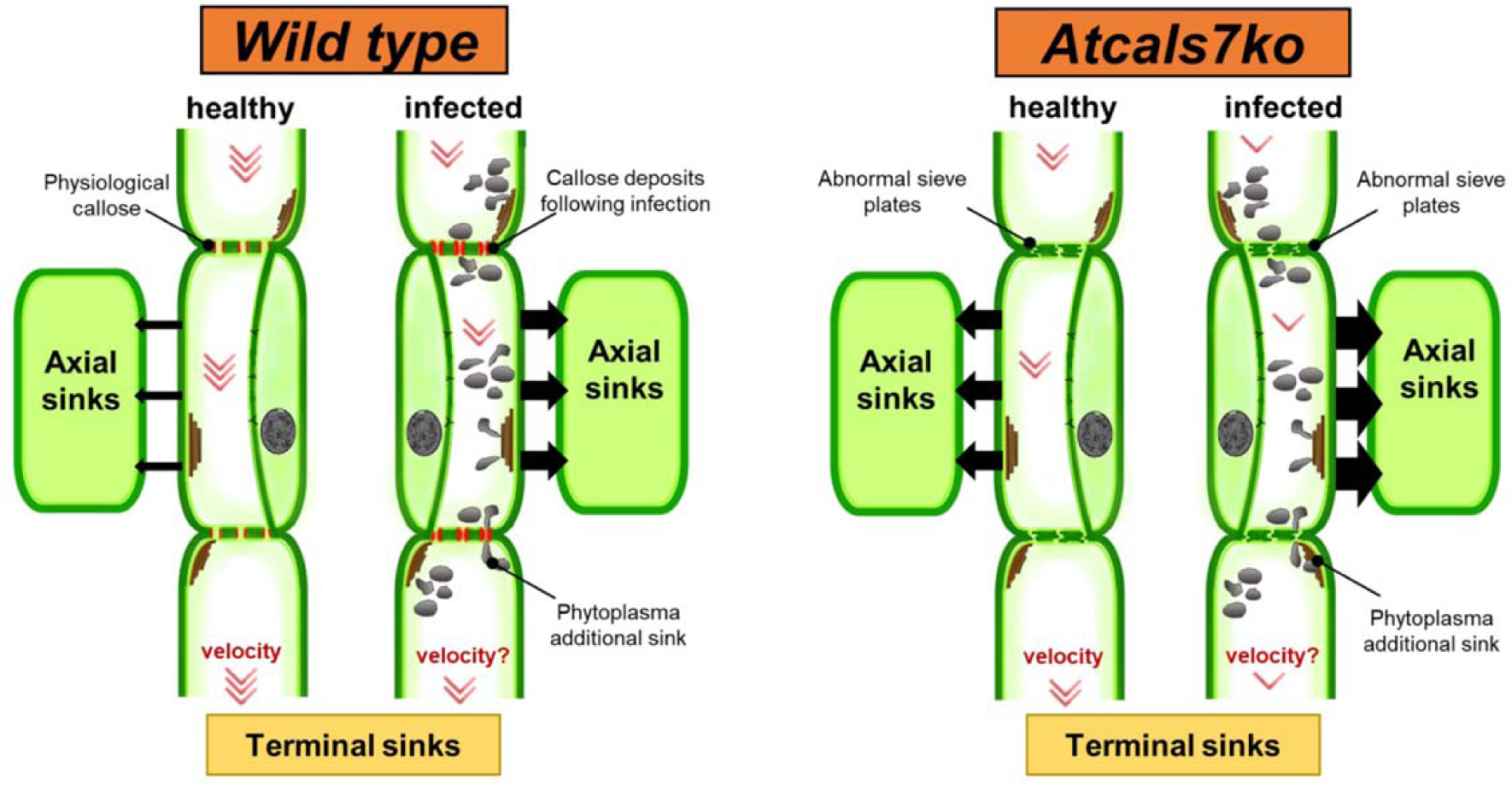
Hypothetical model for the impact of functional (wild-type line) or aberrant (*Atcals7ko* line) sieve plates on the photoassimilate investment into terminal or axial sinks. The red arrowheads quantify the extent of investment in terminal sinks, which is correlated to the translocation speed. The size of the horizontal black arrows quantifies photoassimilates which are invested in axial sinks. Photoassimilate investment into terminal sinks is lower in the *Atcals7ko* line, due to aberrant sieve plates, which results in more escape of carbohydrates along the pathway towards the axial sinks. In case of phytoplasma infection, photoassimilate investment into terminal sinks could be more affected, favoring not only the axial sink proliferation, but also the phytoplasma (additional sink) nourishment.

*AtSUS5* and *AtSUS6* encode two sucrose synthases located in the SEs, which provide UDP-glucose as substrate for AtCALS7. Comparing the expression levels found in healthy samples (i.e. wild-type *versus Atcals7ko*), *AtSUS6* was significantly up-regulated in the mutant line (Fig. 7B). *AtSUS5* showed low expression levels which did not differ in the two lines. Following CY-infection, AtSUS5 transcripts increased 3.5 and 5 times in wild- type and mutant lines, respectively, while AtSUS6 transcripts increased 13.5 and 7.5 times (Fig. 7B). AtCWINV1 and AtCWINV6 encode two cell wall invertases, involved in the irreversible cleavage of sucrose into glucose and fructose in the apoplast. Cell wall invertases represent a path for hexose production, as an alternative to sucrose synthases. The expression level of *AtCWINV1* was significantly higher in the midrib of healthy *Atcals7ko* plants (Fig. 7C) than in the other plant groups. *AtCWINV6* showed a similar trend, but the very low expression level prevented us from calculating a significant difference between the lines (Fig. 7C). The gene encoding the sugar efflux facilitator AtSWEET11 was down-regulated in healthy *Atcals7ko* midribs in comparison with the healthy wild-type (Fig. 7D). In response to infection, it was significantly over-expressed in *Atcals7ko* line (Fig. 7D). Transcription levels of *AtSWEET12* were unchanged in all tested samples (Fig. 7D). The expression level of *AtSTP13*, a gene encoding a vascular hexose transporter, was similar in the midribs of both lines, but was enhanced significantly (around 5.5-fold) in the infected *Atcals7ko* line (Fig. 7E). The transcripts of the two genes *AtSUC2* and *AtSUC3*, encoding sucrose transporters located respectively in CCs and in SEs (Meyer et al., 2004) were analyzed in the midribs of both lines (Fig. 7F). The expression level of *AtSUC2* did not significantly change in all plant groups, while *AtSUC3* was significantly up-regulated only in the *Atcals7ko* line following CY infection (Fig. 7F).

All in all, three general trends were visible regarding the gene expression in infected mutant and wild-type plants in comparison with their healthy controls. Modulation was absent for *AtCWINV1* and *AtCWINV6* in both lines, *AtSUS5* and *AtSUS6* were upregulated in both lines, while *AtSWEET11*, *AtSUC3*, and *AtSTP13* were upregulated in mutants only.

## Discussion

Early reports already noted that sieve pores are rich in callose (Eschrich, 1956), which is produced by transmembrane callose synthases, a gene family that comprises 12 members in Arabidopsis and is responsible for cell- or tissue-type specific callose production (Verma and Hong, 2001). Barratt et al., (2011) and Xie et al., (2011) demonstrated that, in *Arabidopsis thaliana*, callose deposition at the sieve plates is regulated by a callose synthase gene, *AtCALS7*, which is spatially associated with the phloem-specific sucrose synthase genes (*AtSUS*) 5 and 6 (Barratt et al., 2009). The same authors attributed a role to both sucrose synthases in providing the UDP-glucose for callose synthesis around sieve pores. Callose synthesis requires several steps, including the glucosyl-group transfer by a transferase (Hong et al., 2001; Bonke et al., 2003; Barratt et al., 2009), which renders credibility to a callose synthase complex, composed of diverse proteins (Schneider et al., 2016).

For plants infected by phytoplasmas, increased callose deposition at the sieve plates has been described since the ‘70s (Braun and Sinclair, 1978) as a means of constricting sieve pores and limiting pathogen spread (Musetti et al., 2013). Enhanced callose deposition was associated with an upregulation of callose synthase genes in phytoplasma-infected apple tree, grapevine and tomato (Musetti et al., 2010; Santi et al., 2013b; De Marco et al., 2016). The structural and physiological modifications were seemingly consistent with transcriptional changes in sucrose transport and metabolism following infection (Santi et al., 2013b), and appeared worth investigating. As phytoplasmas are phloem-limited pathogens (van Bel and Musetti, 2019), we used healthy and phytoplasma-infected Arabidopsis lines, wild-type or *Atcals7* knock-out (Barratt et al., 2011; Xie et al., 2011) to investigate whether and to what extent the sieve-element specific callose synthase 7 (AtCALS7) interferes with plant sugar translocation and metabolism in response to phytoplasma infection.

### Lack of AtCALS7 affects carbohydrate translocation speed and plant growth

Formulae for mass flow determination (De Schepper et al., 2013 and literature therein) predict that the shape of sieve plates controls the translocation velocity. We measured the speed of carbohydrate transport in wild-type and *Atcals7ko* Arabidopsis plants using a non-invasive method (Vincent et al., 2019). As expected, *Atcals7ko* mutants display a decreased translocation speed consistent with the distorted shape of the sieve pores (Figs. 3J and K and Barratt et al., 2011). Reduction of the linear translocation speed seems to have an effect on the carbohydrate supply of the sinks. As a result, growth is slowed down as expressed by the reduction of rosette fresh weight and stem length in mutant plants (Fig. 2B and C).

The decreased growth indicates that the balance between the supply of terminal and axial sinks is appreciably disturbed in mutants (e.g. Hafke et al., 2005). Since sugar concentrations and expression levels of sugar-handling enzymes are similar in wild-type and mutant plants (Figs. 6 and 7), the disturbance of nutrient distribution is likely due to mechanical alterations. At the same phloem-loading rates, the concentration of soluble carbohydrates must be higher in sieve-tube saps transported at lower velocities. At reduced velocities, moreover, solute retrieval rates by axial sinks along the transport pathway are expected to be higher due to the longer residence times of translocate in the transport pathway (Horwitz, 1958). Both factors effectuate an increased retrieval by axial sinks, even if their carbohydrate transporters are not upregulated (Fig. 8). And finally, a more efficient symplasmic connectivity in the cortex of mutants allows a quicker transfer to and storage by axial sinks (Fig. 4J). Knocking-out AtCALS7 thus restricts the nutrient supply of terminal sinks (Fig. 8), while the axial sinks seem to profit from the reduced translocation speed (Fig. 1).

Phytoplasma infection highlights another mode of AtCALS7 impact. In response to CY infection, mutants and wild-type plants showed respectively an additional 27% and 16% rosette-weight loss in comparison with their healthy controls. CY-infected plants were incapable of developing normal stems (Fig. 2C and D), with mutant plants showing stronger effects. Even if the increased callose deposition did not constrict all sieve plates (Fig. 3F and G; Braun and Sinclair, 1978; Gallinger et al., 2021), stem stunting reported in CY-infected wild-type plants (Pagliari et al., 2016) is indicative of a strong disturbance of phloem functions imposed by phytoplasmas (Maust et al., 2003). The modified distribution patterns suggest that nutrients are invested in additional sinks in the form of phytoplasmas at the cost of the terminal sinks (Fig. 8). The investment in phytoplasma growth would be in the mutant plants twice as high as in the wild-type plants. As a matter of fact, the phytoplasma titre is indeed about twice as high in the mutants (Fig. 2E). Phytoplasmas seem to exploit the absence of At*CALS7* for better recruitment of resources for their growth. Below, we will explore in the sections the underlying factors that explain the higher fitness of phytoplasmas in mutant plants.

### The altered symplasmic communication between cortex cells in the *Atcals7ko* line favors phytoplasma infection

Plasmodesmata and their gating mechanisms could be of importance for the survival, dissemination and effector dispersal of phytoplasmas (van Bel and Musetti, 2019 and literature therein). In this context, callose deposition that influences plasmodesmal functional dynamics, by modulating the passage of molecules through the cytoplasmic sleeve, is of paramount importance (Sager and Lee, 2018).

To check the callose distribution and deposition level in different tissues of wild-type and *Atcals7ko* Arabidopsis midribs, healthy or CY-infected, aniline-blue that becomes fluorescent by binding to callose, was used (Zavaliev and Epel, 2015). As expected, the fluorescent dots appeared in the SE/CC area in the midribs of wild-type plants (De Marco et al., 2016; Pagliari et al., 2016) but not in the *Atcals7ko* mutants. This is in agreement with the TEM results (Fig. 3; Barratt et al., 2011; Xie et al., 2011). In midrib cortical parenchyma of both Arabidopsis lines, aniline-blue staining showed callose deposition at the plasmodesmata, as numerous fluorescent punctate dots at the cell boundaries (Levy et al., 2007). In the two Arabidopsis lines, the number of deposits and the fluorescence intensity was not significantly different, in average, within the region of interest. Nevertheless, considering the integrated density of the dots in that area, this was significantly higher in the *Atcals7ko* line than in the wild-type, indicating differences in plasmodesmal functional dynamics. It has been recently suggested that callose could increase the elasticity rather than rigidity of the wall matrix (Abou-Saleh et al., 2018), favoring selective symplasmic transport. This is in line with a more efficient symplasmic connectivity in the cortex of mutants to promote axial sink nourishment, as suggested above.

Callose level was strongly enhanced in cortical tissues of infected wild-type plants, but not in the mutant (Fig. 4H). This lack of additional callose deposition suggests that the cortical plasmodesmata in mutants do not respond to phytoplasma infection. This may have two consequences in mutant plants. First, effectors may disseminate more effectively into extra-fascicular tissues (van Bel and Musetti, 2019) and second, hexose and sucrose move more easily from cell to cell.

In the epidermal tissue, the number of fluorescent dots, the signal intensity and the integrated density is lower in *Atcals7ko* samples compared to the wild-type, suggest an attempt of mutant plants to establish the symplasmic continuity in tissues far from the phloem, to control symplasmic return of assimilates towards the region adjoining the phloem. In case of phytoplasma infection, callose deposition at epidermis level increases in the mutant line, which infers that plasmodesmata increase the capability to control symplasmic molecular movement also in tissues not directly involved in pathogen interaction.

### Phytoplasma infection increases sugar contents in midribs of *Atcals7ko* plants

In midribs of healthy source leaves, sugar levels (i.e. sucrose, glucose, fructose, myoinositol, sorbitol, raffinose, arabinose) were similar in the wild-type and *Atcals7ko* Arabidopsis lines. The production of sucrose and its related catabolites is not influenced by the loss of AtCALS7, although the exact metabolite distribution per cell type is uncertain due to the use of entire midribs with diverse cell types.

An enhanced rate of carbohydrate metabolism in response to phytoplasma infection is derived from increased glucose, sucrose and myoinositol levels in midribs of mutant and wild-type plants (Santi et al., 2013a, 2013b; Yao et al., 2019). This upregulation of the soluble sugar level is more significant in mutants than in wild-type plants (Fig. 7). In the wild-type line, the significantly increased level of *AtSUS5* and *AtSUS6* transcripts (Fig. 7) is in line with the higher expression level of *AtCALS7* (Fig. 7; Barratt et al., 2011), to provide precursors (i.e. UDP-glucose) for callose synthesis (Tan et al., 2015; De Marco et al., 2020), and for the production of carbon skeletons for synthesis of defense-related compounds (Bolouri-Moghaddam et al., 2010; Musetti et al., 2013). In *Atcals7ko* infected plants, increased photoassimilate investment into axial and additional sinks (such as phytoplasmas) would explain enhanced carbohydrate content and the up-regulation of *AtSUS5* and *AtSUS6* genes, which provide energy at the infection site (Yao et al., 2019; 2020). In the infected mutant line high amount of myoinositol was detected. In the absence of callose, myoinositol could be useful to modulate SE wall plasticity as an attempt to respond to the stress imposed by phytoplasmas, as demonstrated for other environmental cues (Wu et al., 2018). Myoinositol is indeed an excellent carbon source for cell wall constituents, such as pectin and hemicellulose (Loewus et al.,1962), which are, together with cellulose, the major types of polysaccharides in Arabidopsis cell walls (Bethke et al., 2016).

### Loss of *AtCALS7* leads to concerted channeling of photoassimilates to CY-infected sieve tubes

The question now arises, if the sugar upsurge is due to a higher activity of enzymes involved in oligosaccharide metabolism and/or to upregulation of transporters directing these substances to the SE/CC complexes in the midribs.

Increased levels of SE-restricted *AtSUS5* and *AtSUS6* expression (Fig. 7) suggest an enhanced need for sugar processing in SEs. Since CWINV1 and CWINV6 are not upregulated in infected mutant and wild-type plants (Fig. 7), apoplasmic breakdown of sucrose is likely not critical for the sugar supply of phytoplasmas. By contrast, sugar transport capacity to phytoplasmas (and thus to SEs) seems to be limited in wild-type plants. Without exception, transporters involved in sugar uptake by SE/CC complexes are upregulated in healthy mutant plants (Fig. 7), but not in wild-type plants. The higher expression rate, probably subject to effector control, renders it plausible that the better symplasmic mobility of phytoplasma effectors (Fig. 5) make them more effective in mutants.

*AtSUC2* and *AtSUC3*, encode for transmembrane sucrose cotransporters localizing respectively in CCs and in SEs (Meyer et al., 2004). AtSUC2 and AtSUC3 are not only responsible for the sucrose loading, but are also involved in the sucrose retrieval along the transport phloem (Meyer et al., 2004; Gould et al., 2012).

*AtSTP13* belongs to a family of genes encoding for hexose cotransporters. *AtSTP13* is strongly expressed in leaf main veins (Yamada et al., 2011), where it contributes to the basal resistance to necrotrophic fungi (i.e. *Botrytis cinerea*, Lemonnier et al., 2014) and extracellular bacteria (i. e. *Pseudomonas syringae*, Nørholm et al., 2006) by depriving these pathogens of modified sugar fluxes toward host cells. On the contrary, the over- expression of *STP13* in plants infected with intracellular biotrophic pathogens provokes increased host susceptibility (Huai et al., 2020), by promoting cytoplasmic hexose accumulation. As phytoplasmas are biotrophic intracellular pathogens, it is conceivable that the over-expression of *AtSTP13* in CY-infected *Atcals7ko* plants could also indicate an increased susceptibility of this line to phytoplasma attack, in comparison to the wild-type (Bezrutczyk et al., 2018). Indeed, *Atcals7ko* line appears more affected by infection (i.e. Fig. 2).

As for AtSUCs, AtSWEET11 and AtSTP13 are also involved in the retrieval of sucrose along the transport phloem (Meyer et al., 2004; Gould et al., 2012). Taken together, the significant increase of *AtSWEET11*, *AtSUC3* and *AtSTP13* transcripts in the midribs of infected *Atcals7ko* plants indicates that sugar retrieval along the phloem path is highly activated in the host to fuel pathogen proliferation (Tadege et al., 1998). While AtSUC2, AtSUC3, and AtSTP3 are localized to the SE/CC complexes (Fig. 9), the cellular deployment of SWEET11 and SWEET12 is less certain. It must be underscored that midribs contain SE/CC complexes with a high degree of release and retrieval due to symplasmic isolation under source-limiting conditions. If SWEETs are responsible for sucrose supply of phytoplasma, they may mediate import at the SE/CC complex plasma membrane or export at the phloem parenchyma plasma membrane (Fig. 9).

**Fig. 9.**
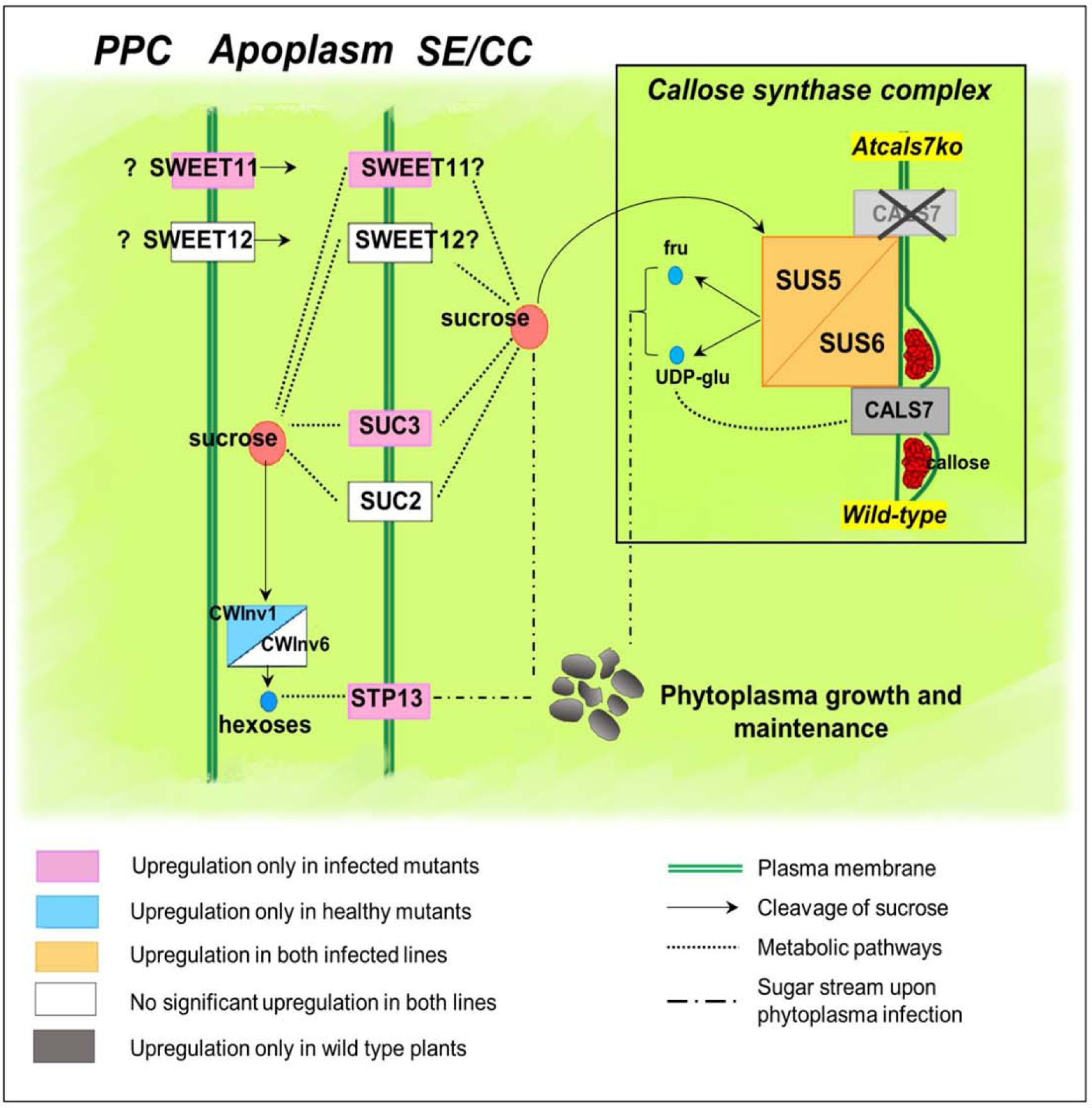
Localization of the proteins involved in sugar metabolism and transport in the midribs of wild- type and *Atcals7ko* Arabidopsis and their expression in response to phytoplasma infection. In transport phloem, AtSUC2, AtSUC3, and AtSTP13 are localized to the plasma membrane of SE/CC complexes, while deployment of SWEET11 and SWEET12 is less certain, as they localized to the phloem parenchyma (Le Hir et al., 2015) but also to the companion cells (Abelenda et al., 2019). AtSUS5 and 6 are localized to the SE plasma membrane. Independently from the different protein localization, and regardless of the carbohydrate form (sucrose or hexoses), up-regulation of *AtSUC3*, *AtSTP13* and *AtSWEET11* in *Atcals7ko* plants would stimulate sucrose transport to SE/CC complexes (dotted lines), supporting phytoplasma maintenance (dashed lines) and giving rise to an increased susceptibility of the mutant line to phytoplasma infection.

*AtSWEET11* and *12* were characterized as sucrose effluxers localized to the plasma membrane of phloem parenchyma cells with a role in feeding active transport into the SE/CC complex in the collection phloem (Chen et al., 2012). They are expressed in most *Arabidopsis* tissues (Braun, 2012), so, beyond phloem loading, they might play other roles (Breia et al., 2021). In the current understanding, deployment of AtSWEET11 and AtSWEET12 in phloem parenchyma of floral stems is quite meaningful (Le Hir et al., 2015). OsSWEETs are also tentatively localized to phloem parenchyma in the phloem-unloading zone of rice (Milne et al., 2018), but StSWEET11 was proposed to be deployed in the plasma membrane of companion cells of potato transport phloem (Abelenda et al., 2019). With the present arguments in hand, it seems plausible that SWEET11 and SWEET12 are localized to the phloem parenchyma (Fig. 9).

The possibility that SWEETs determine plant susceptibility or resistance by the control of nutrient provision to pathogens has recently been discussed (Chen et al., 2010; Bezrutczyk et al. 2017; Breia et al., 2021). *AtSWEET11* is a member of the clade III of the SWEET genes, involved in disease development (Li et al., 2018). Wipf et al., (2021) reported that AtSWEET11 interacts with AtRBOHD, a membrane NADPH oxidase producing reactive oxygen species, involved in defense-related processes. The constitutive downregulation of *AtSWEET11* in the mutant line could be related to a less effective capability to react to pathogen attacks. In this regard, the increased expression level of *AtSWEET11* in infected mutant plants (with the simultaneous over expression of *AtSUC3*) could promote sucrose release towards the SE/CC complexes to support phytoplasma nourishment and proliferation (Fig. 9).

### Concluding remarks

At*CALS7* appears to have two layers of action as demonstrated by *Atcals7ko* mutants. As a first layer of action, knocking out *AtCALS7* strongly restricts the supply of terminal sinks with resources and enhances the supply of axial sinks (Fig. 8). This results in a severely reduced growth of host plants. The second layer of action becomes manifest by an increased inability to counter infection due to an increased symplasmic connectivity in *Atcals7ko* mutants, which allows a better spread of phytoplasma effectors. The effectors probably stimulate the expression of genes involved in channeling resources towards the phytoplasmas residing in sieve tubes (Fig. 9). The enhanced withdrawal of sugar reserves from the host metabolism favours phytoplasma growth and an additional plant growth reduction in *Atcals7ko* mutants.

## Materials and methods

### Plants and insect vectors

Seeds of wild-type and Atcals7ko (SALK_048921) lines of *Arabidopsis thaliana* ecotype Col0 were provided by the Nottingham Arabidopsis Stock Centre (NASC). Sixtyfour plants were grown in a climate chamber for 40 days on 5:1 mixture of soil substrate and perlite and fertilized twice a month with an N-P-K liquid fertilizer, as described previously (Pagliari et al., 2017), 32 plants under short-day light conditions (9h L/15h D) at 18-20°C, and 32 under long-day light conditions (14h L/10h D), at 23°C in order to develop the floral stem for the transport speed experiment.

Healthy colonies of *Macrosteles quadripunctulatus* were reared on *Avena sativa* in vented plexiglass cages in greenhouse (temperature of 20-22°C and short-day conditions 9h L/15h D), according to Bosco et al. (1997). The last instar nymphs were transferred to Chrysanthemum carinatum plants infected with Chrysanthemum Yellows (CY) phytoplasma (Lee et al., 2003), a strain related to ‘*Candidatus* Phytoplasma asteris’ (‘*Ca*. P. asteris’, 16SrI-B subgroup), as the source of inoculum for a 7-day phytoplasma acquisition-access period (AAP). After a latent period (LP) of 18 days, insects were transferred to 40-day-old Arabidopsis plants for the 7-day inoculation-access period (IAP).

Eight wild-type and eight *Atcals7ko* plants were exposed to 3 infectious *M. quadripunctulatus* individuals (CY-infected plants). In addition, sixteen plants from both lines (8+8) were exposed to 3 healthy vectors as a control (healthy plants). Healthy leafhoppers were collected from healthy colonies and were of the same age as the infected ones. When symptoms on the infected plants grown under short-day conditions were clearly visible, i.e. 26 days post inoculation (Pacifico et al., 2015; Pagliari et al., 2017), midribs from the leaves of the third rosette node (source leaves) were collected. For phytoplasma detection and gene expression analyses, collected samples were immediately frozen in liquid nitrogen and stored at -80°C until use. For carbohydrate analyses, midribs were immediately frozen in liquid nitrogen, freeze-dried and then stored at -80°C until use.

### Phloem transport speed measurement

For phloem transport experiments, Arabidopsis plants were grown as reported above, under long-day light conditions (14h L/10h D), at 23°C. Sixty-day-old healthy plants of both lines, were used for the measurement. Two x-ray photomultiplier tubes, 5.6 cm in diameter, (St. Gobain, Malvern, PA) were placed along the floral stem of both wild-type and *Atcals7ko* plants, leaving a 2-cm-long buffer zone between the detectors, while a third detector monitored the signal from the rosette.

Each detector was connected to an M4612 12-channel counter and counts of x-rays were logged with the manufacturer’s software (Ludlum Measurements, Sweetwater, TX, USA). Plants were placed on a lead layer having a receptacle, under the same day length used for growing. After at least 8 hours of background measurement, 600 μl of NaH^14^CO_3_ solution (specific activity: 40–60 mCi (1.48–2.22 GBq)/mmol) were injected into the receptacle. Immediately thereafter, 1000 µl of a saturated citric acid solution was also injected into the receptacle. Plants were tightly closed in a plastic bag, and were allowed to assimilate ^14^CO_2_ for 2 hours. Then the bag was removed and the remaining gaseous radiolabelled isotope was flushed *via* the fume hood. The x-ray detectors monitored the x-ray counts from the plant tissue every minute for 72 hours. The distance between the detectors was measured at the end of the experiment.

The data of the counts from each detector were handled with the *<NMLE>* package in Rstudio (Pinheiro et al., 2018). In accordance with previously published methods, the Rstudio script was written to create a logistic function, and to fit the logistic function to the data with time on the x-axis and the recorded counts on the y-axis (Vincent et al., 2019). Xmid, which represents the time at the half-height of the logistic curve or the average time of arrival of ^14^C in the phloem tissue near the x-ray detector, was calculated for each detector placed along the stem. The speed of translocation was calculated by dividing the distance between the middle of the two detectors (in cm) and the difference between the Xmid timepoints from the detectors (in h). Both infected lines failed to normally develop flowering stalks: stalk was very short in infected wild-type plants and absent in infected *Atcals7ko* (Fig. 2C). For these reasons, it was not possible to calculate and compare sugar translocation speed in the two infected Arabidopsis lines. For both wild-type and *Atcals7ko* healthy plants 4 measurements were carried out, pairing 1 wild-type and 1 *Atcals7ko* together in each labelling.

### Symptom description

Symptom development was observed in healthy and CY-infected plants of both lines, from the end of the inoculation period to the time of tissue harvest for different analyses, i.e. when plants were ca. 70 days old. To evaluate the phenotypic differences between the two lines and the differences due to phytoplasma infection, the rosette fresh weight was measured in plants grown under short-day conditions using 10 healthy and 10 CY-infected plants per line. The day before sampling, soil was saturated with water. Rosettes were then cut at the plant collar level and the fresh weight of each biological replicate was measured. Moreover, the length of the floral stalk was measured in plants grown under long-day light conditions (see above), using 8 healthy and 8 CY-infected plants per line. Statistical analyses were performed using RStudio software Version 1.1.456 (RStudio Team 2020, Boston, MA). The normal distribution was checked with the Shapiro-Wilk test. Significant differences among the group means were determined with a two-way ANOVA and post-hoc comparisons between all groups were made with Tukey’s test with P<0.05.

### Phytoplasma detection and quantification

To check the phytoplasma titre in wild-type and *Atcals7ko* CY-infected plants, genomic DNA was extracted from 200 mg of fresh leaf midrib tissue, according to Doyle and Doyle protocol, (1990) modified by Martini et al., (2009). The ribosomal protein gene *rplV* (*rpl22*) was chosen as a target for the amplification of CY phytoplasma DNA using the primer pair rp(I-B)F2/rp(I-B)R2 (Lee et al., 2003; Pagliari et al., 2017) and a CFX96 real-time PCR detection system (Bio-Rad Laboratories, Richmond, CA, USA). A standard curve was established by 10-fold serial dilutions of a plasmid DNA containing the 1260 bp ribosomal protein fragment from CY phytoplasma, amplified with the primer pair rpF1C/rp(I)R1A (Martini et al., 2007). Real-time PCR mixture and cycling conditions were previously described (Pagliari et al., 2017). The phytoplasma titre was expressed as the number of CY-phytoplasma genome units (GUs) per mg of fresh leaf sample to normalize the data. Statistical significance of the quantitative differences between phytoplasma populations was calculated by analysis of three replicates of 8 plants per line. Statistical analyses were performed using RStudio software version 1.1.456 (2009-2018 RStudio, Inc. Boston, MA). The normal distribution was checked with Shapiro-Wilk test. Significant differences among the means were determined using the Kruskal-Wallis non-parametric test with P<0.05.

### Gene expression analyses

Total RNA was extracted from 100 mg of leaf midrib powder, from 5 plants for the four experimental conditions, obtained by grinding in liquid nitrogen and using a SpectrumTM Plant Total RNA kit (Sigma-Aldrich, Merck KGaA, Darmstadt, DE) according to the manufacturer’s instructions. The RNA reverse-transcription was carried out using a QuantiTectReverse Transcription Kit (Qiagen N.V., Hilden, DE) following the manufacturer’s instructions. The expression of the genes, reported in Table 1, was analysed in healthy and CY-infected midribs by real-time experiments, performed on a CFX96 real-time PCR detection system (Bio-Rad Laboratories). The reference gene was selected by comparing the *AtUBC9* (ubiquitin conjugating enzyme 9), *AtTIP41* (TIP41-like family protein), *AtSAND* (SAND family protein), and *AtUBQ10* (polyubiquitin 10) gene expression. The gene stability values (M values) were calculated according to the geNorm program (Pagliari et al., 2017). *AtUBC9* gene was the most stably expressed gene and so the most suitable as a reference gene (M=0.44). SsoFast EvaGreen Supermix 2x (Bio-Rad Laboratories Inc., Hercules, CA, USA) and cDNA obtained from 5 ng of RNA and specific primers (Table1) were used in a total volume of 10 μl. Under these conditions, the primer pair efficiency was evaluated as described by Pfaffl (Pfaffl, 2001) using standard curves of different dilutions of pooled cDNA. PCR was performed as described in Pagliari et al., (2017), with three technical repeats. A mean normalized expression (MNE) for each gene of interest (Muller et al., 2002) was calculated by normalizing its mean expression level to the level of the UBC9 gene. Five individuals were used for the gene MNE determination. Statistical analyses were performed using RStudio software Version 1.1.456 (2009-2018 RStudio, Inc. Boston, MA). The normal distribution was checked with Shapiro-Wilk test. Significant differences among the means were determined by a two-way ANOVA and post-hoc comparisons between all groups were made with Tukey’s test with P<0.05.

**Table 1:**
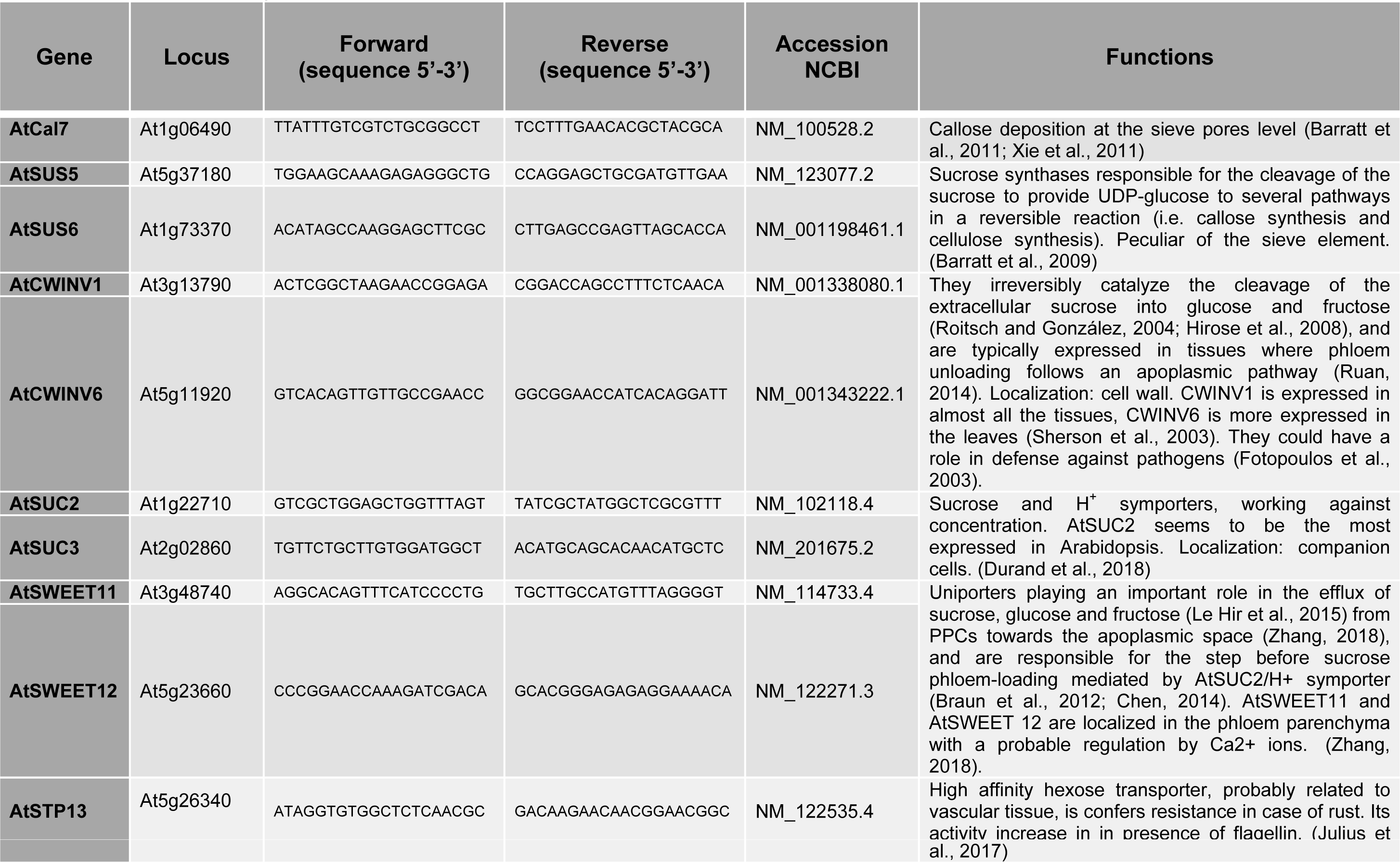
List of the genes analyzed in this work. For every gene the locus, the primer sequences, the accession number (NCBI)and the function has been reported.

### Transmission electron microscopy

To observe phloem ultrastructure in the midrib, samples were prepared for microscopic analyses as reported by Pagliari et al., (2016). For each condition, at least five segments (10 mm in length) of the mature leaf midrib from 5 individuals were submerged in MES buffer (10mM NaOH-2-(*N*-morpholino) ethanesulfonic acid, 2mM CaCl_2_, 1mM MgCl_2_, 0.5mM KCl and 200mM mannitol, pH 5.7) for 2h at room temperature. Samples were fixed in a solution of 3% paraformaldehyde and 4% glutaraldehyde for 6 h, the solution was refreshed every 30 minutes. After rinsing, samples were post-fixed overnight with 2% (w/v) OsO_4_, dehydrated with an ethanol gradient and transferred to pure propylene oxide. Midrib segments were then embedded in Epon/Araldite epoxy resin (Electron Microscopy Sciences, Fort Washington, PA, USA). Ultra-thin sections were collected on uncoated copper grids, stained with UAR-EMS (uranyl acetate replacement stain, Electron Microscopy Sciences) and then observed under a PHILIPS CM 10 (FEI, Eindhoven, The Netherlands) transmission electron microscope (TEM), operated at 80 kV, and equipped with a Megaview G3 CCD camera (EMSIS GmbH, Münster, Germany). Five non-serial cross-sections from each sample were analysed.

### Confocal laser scanning microscopy

In both plant groups, healthy or CY-infected, callose deposits, associated with plasmodesmata or sieve plates, were identified in leaf epidermis, midrib cortical parenchyma, and SE/CC areas, using aniline blue staining and Confocal Laser Scanning Microscopy (CLSM). Callose was quantified *in situ* by measuring the aniline blue fluorescence intensity (Levy et al., 2007). For analysis of the epidermis, whole leaves from the third rosette node were collected in 95% EtOH solution and stored for at least 2 hours at room temperature, as described by Zavaliev and Epel (2015). Samples were rehydrated with distilled water with 0.01% Tween-20 for 1 hour, then placed in a tube with aniline blue solution (0.01M K_3_PO_4_, pH=12). Tubes with samples were placed in a vacuum desiccator for 10 minutes, then incubated under aluminum foil for 2 hours. Two leaves from three plants for every condition (i.e. wild-type or *Atcals7ko*, healthy or CY-infected) were observed. For every leaf at least 20 images were taken on a single-plane, using a Leica SP8 LSCM (Leica Microsystems Inc., Buffalo Grove, IL, USA) equipped with 40x oil immersion objective. Aniline blue fluorescence was excited with 405nm diode laser and emission was detected at 475-525nm. Optimal conditions for microcopy were previously determined using healthy wild-type samples.

To analyze callose deposits in the midribs, both in the SE/CC area and in the surrounding midrib cortical parenchyma, leaf midribs were collected in a 95% EtOH solution and stored at least 2h at room temperature. Hand-made cross sections, cut with a razor blade, were incubated for 5 minutes in aniline blue solution. The sections were rinsed with 0.01% Tween-20 solution. For each experimental condition at least 10 non-serial sections from 5 different specimens were observed using a Leica SP8 CLSM with a 10x objective. Aniline blue fluorescence was excited with a 405 nm diode laser and emission was detected at 475-525 nm. Optimal conditions for microscopy were previously determined by using healthy wild-type samples.

Image analysis was performed using FIJI software (Schindelin et al., 2012). For the analyses of epidermis and midrib cortical parenchyma, images were treated following the protocol by Zavaliev and Epel, (2015). A macro separated the fluorescent signal in the region of interest from the background using an auto local threshold with the Phansalkar algorithm with radius = 1 for epidermal tissue, and the Bernsen algorithm with radius = 10 for midrib cortical parenchyma. For cortical parenchyma, an area of interest of 45000 μm^2^ was analyzed.

Fluorescent dots (indicating callose deposits) in the SE/CC area were analyzed as previously reported by Pagliari et al., (2016). For every group of samples, the mean gray value (callose intensity), the number of callose deposits for each square millimeter and the integrated density (sum of intensity within the area of the region of interest) were considered. Data obtained by analyses were handled using RStudio software (2009-2018 RStudio, Inc. Boston, MA). Normality of the data was checked with the Shapiro-Wilk test, outliers were removed according to the RStudio function (boxplot(data$variable, plot=FALSE)$out), and data were normalized, where necessary, with Box-Cox transformation. A two-way analysis was performed followed by a post-hoc comparison of all pairs with Tukey’s test, with P<0.05.

### Sugar quantification

Authentic standards of sugars (rhamnose, arabinose, fructose, glucose, maltose, sucrose, and melibiose) and sugar alcohols (glycerol, myoinositol, arabitol, and sorbitol) were purchased from Sigma-Aldrich (St. Louis, MO, USA). Internal standards including fructose-^13^C6 (for sugars) and sorbitol-^13^C6 (for sugar alcohols) were obtained from Toronto Research Chemicals (Toronto, ON, Canada) and Sigma-Aldrich (St. Louis, MO, USA), respectively. Water, acetonitrile, methanol, and formic acid were of LC-MS grade, and were purchased from Fisher Scientific (Fair Lawn, NJ, USA). Stock solutions of each analyte and internal standard were prepared at a concentration of 10,000 μg/ml in water, or methanol. Working standard solutions were prepared by diluting and mixing each stock solutions with 90% methanol (water/methanol = 10/90, v/v). The stock and working solutions were stored at -80°C. Freeze-dried leaf midribs were stored at -80°C until use. Ten milligram of ground samples treated with 0.05 ml of internal standard solution (50 μg/ml sorbitol-13C6 and 200 μg/ml fructose-13C6 in 90% acetonitrile (water/acetonitrile = 10/90, v/v) was extracted with 0.95 ml of 90% acetonitrile (water/acetonitrile = 10/90, v/v) (total volume: 1 ml) by ultra-sonication for 30 min, followed by agitation for 30 min. After centrifugation (20,000 g, 5 min, 4°C), supernatant was further filtered through 0.22 μm nylon filter, and was injected into LC–MS/MS for analysis. The extraction was performed in triplicate using 4 biological replicates. Data obtained by analyses, were handled using RStudio software. Normality of the data was checked with Shapiro-Wilk test, outliers were removed, and data were normalized, where necessary, with Box-Cox transformation. A two-way analysis was performed followed by post-hoc pairwise comparison of all groups with Tukey’s test, with P<0.05.

## Acknowledgements

This work was funded by the University of Udine, through the Department of Agriculture, Food, Environment and Animal Sciences (Di4A), Project Start-up 2018. The authors thank Professor Domenico Bosco (University of Torino, Italy) for making available the CY phytoplasma strain and for his advice on insect rearing.

## Author contribution

CB and RM conceived the project. CB grew the plants, reared the insects and prepared inoculum sources with the help of AL and FB. CB, SB and MM performed the molecular analyses. AmL conceived sugar quantification and the flow speed experiment with the support of CV. CB and SW performed confocal observations. CB, JHS and YW performed the sugar quantification assays. CB with the supervision of MOP performed the flow speed experiment. CB and RM and AJEvB wrote the manuscript with the valuable support of SS. All the authors provided critical suggestions for the realization of the manuscript.

